# Representing cells as sentences enables natural-language processing for single-cell transcriptomics

**DOI:** 10.1101/2022.09.18.508438

**Authors:** Rahul M. Dhodapkar

## Abstract

Gene expression matrices commonly used in single-cell transcriptomics, cannot be directly analyzed with tools developed for natural languages. By restructuring these matrices as abundance-ordered sequences of genes, we generate cell sentences: rank-normalized, positionally encoded expression data. We show that these cell sentences can be analyzed using existing tools from natural language processing to unify cell and gene representations across species.

## 1 Main

Since the first study directly measuring messenger RNA in the individual cells of early stage mouse embryos in 2009, single-cell transcriptomics have been extensively employed to characterize complex biological systems across a wide range of conditions and species [1, 2]. Over the same time period, breakthroughs in natural language processing have made great fundamental contributions to the field of machine learning, including the development of word embeddings and transformer architectures, enabling the training of high-performance models such as GPT-3 [3, 4, 5].

Multiple tools have been developed that apply new approaches from deep learning to the analysis of single cell transcriptomics data for tasks such as batch integration, imputation, denoising, and dimensionality reduction [6]. In all of these approaches, gene expression data is represented by a matrix encoding transcript abundance per cell, obtained from preprocessing tools such as cellranger and STARsolo, after alignment of sequencing reads to an annotated reference [7, 8]. While many general purpose tools already exist in the field of natural language processing (NLP) for a wide variety of tasks, these tools require data represented as strings of words, and are not directly compatible with the raw or normalized expression matrix formats [9].

In this manuscript, we present cell2sentence, a novel method for the transformation of expression matrices to abundance-ordered lists, where genes are analogous to words, and cells are analogous to sentences. Construction of these *cell sentences* enables the direct application of tools from NLP to the analysis of single cell sequencing data. We show that the cell sentence representation captures biologically meaningful information by demonstrating that two commonly used methods from NLP—edit distance and word embedding—can reproduce findings consistent with current standard tools employed for single cell RNA sequencing analysis. We then apply a word embedding alignment procedure, previously developed for cross-lingual translation, to define a new method for the cross-species integration of single-cell transcriptomics data.

The core insight of cell2sentence is to re-organize cell expression matrices into sequences of genes for each cell, ordered by decreasing transcript abundance (Fig 1A). The generation of rank-ordered cell sentences is similar to rank normalization of the gene expression matrix, which has been used previously in gene expression analysis to improve robustness to noise [10]. These cell sentences can be directly rendered as space-delimited text, in a manner similar to natural language.

**Figure 1:**
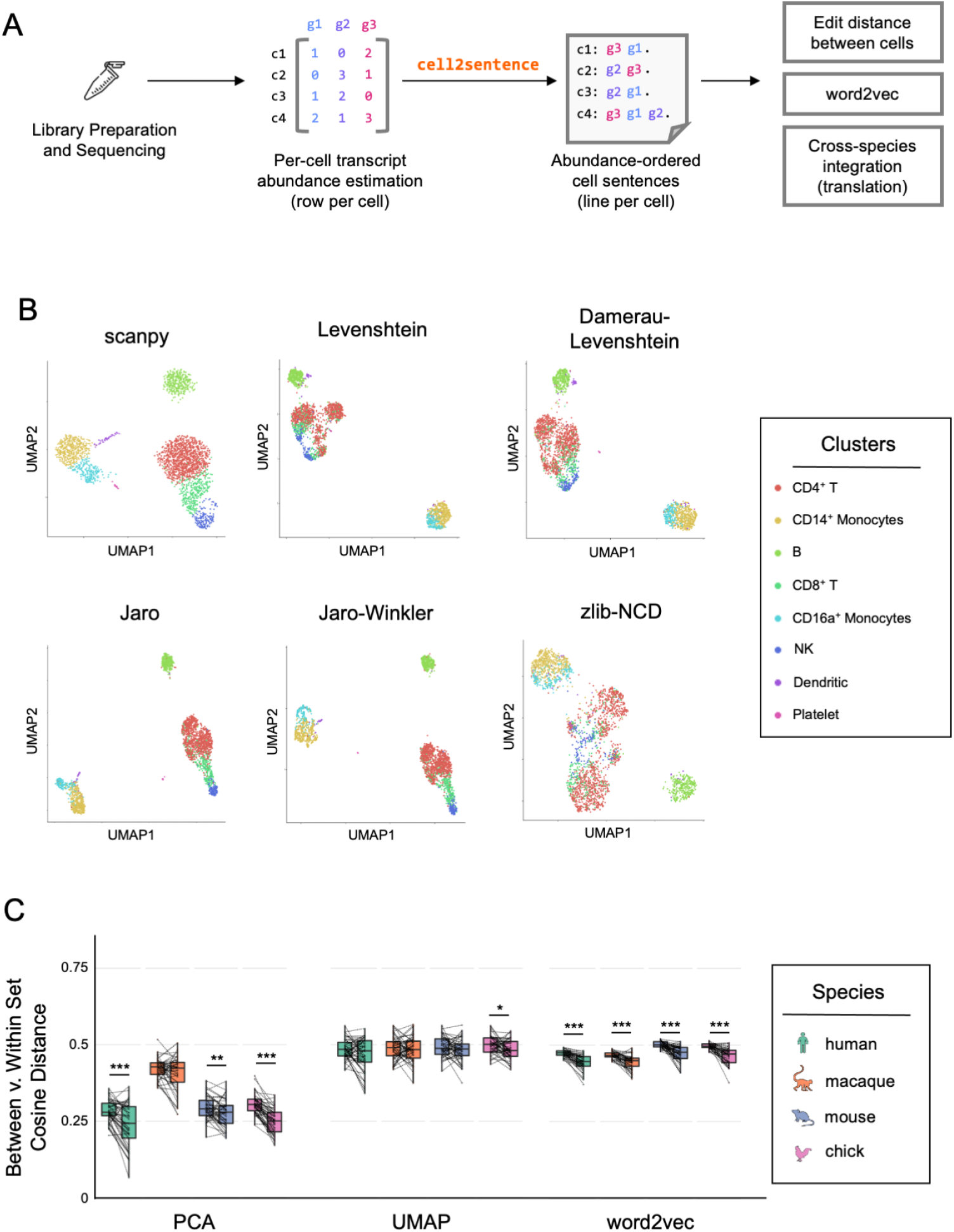
Cell sentences encode biologically meaningful variation, which can be analyzed using tools from natural language processing. (A) cell2sentence transforms gene expression matrices to cell sentences for use with a wide range of downstream applications. (B) Several measures of edit distance (Levenshtein, Damerau-Levenshtein, Jaro, Jaro-Winkler, zlib-normalized compression distance) robustly capture cell types defined using standard methods (scanpy). (C) Gene pairs in the same hallmark gene set are significantly closer in the word2vec gene embedding than other gene pairs. NCD: normalized compression distance, *: p < 0.5, **: p < 0.01, ***: p < 0.001

Edit distance measures the number of operations (or edits) needed to transform one string of symbols to another, and is commonly used to measure the similarity between two sentences [11]. A number of edit distances have been previously described, differing based on the types and costs of allowed transformation operations, as well as the relative positions of mismatches. We generated cell sentences from approximately 3,000 peripheral blood mononuclear cells from a healthy donor, and compared the Levenshtein, Damerau-Levenshtein, Jaro, Jaro-Winkler, and zlib-normalized compression distances against standard expression normalization and distance calculation from the scanpy toolkit [11, 12, 13]. While differences exist between the different methods for distance calculation, all were able to correctly cluster cells annotated using current standard practices (Fig 1B).

Tools such as word2vec use neural models trained on a natural language corpus to map words onto points in an embedded space where position and distance correspond to semantics, termed a word embedding [3]. Other groups have previously generated analogous *gene embeddings* using co-expression patterns, but this required the development of a dedicated training architecture specifically for gene expression data [14]. After using cell2sentence to transform an expression matrix to a set of cell sentences, we are able to directly train the reference implementation of word2vec against single cell transcriptomic data from neural retina of four separate species: human, rhesus macaque, mouse, and chicken [15, 16, 17, 18].

We used hallmark gene sets from MSigDB were used to assess the validity of the word2vec generated embeddings, under the assumption that two genes within a gene set (within set) should be closer in the embedded space than any two arbitrary genes (between set) [19]. We compared the between set and within set cosine distances for gene embeddings generated using word2vec against those derived from principal components analysis (PCA) as well as uniform manifold approximation and projection (UMAP) with matched dimensionality (Fig 1C). While all three methods showed some degree of increased embedding coherence within gene sets, word2vec was the only method that showed significantly reduced within set distance across all species tested (Fig 1C).

To further validate the gene embeddings obtained from word2vec, we generated cell sentence embeddings through mean pooling for each of the four species profiled (Fig 2A-B). Previously published cell type labels obtained from standard analyses were used to color plots of the cell sentence embeddings. These plots demonstrated that for all species analyzed, cell sentence embeddings were able to recover expert-annotated cell types from the sequencing data.

**Figure 2:**
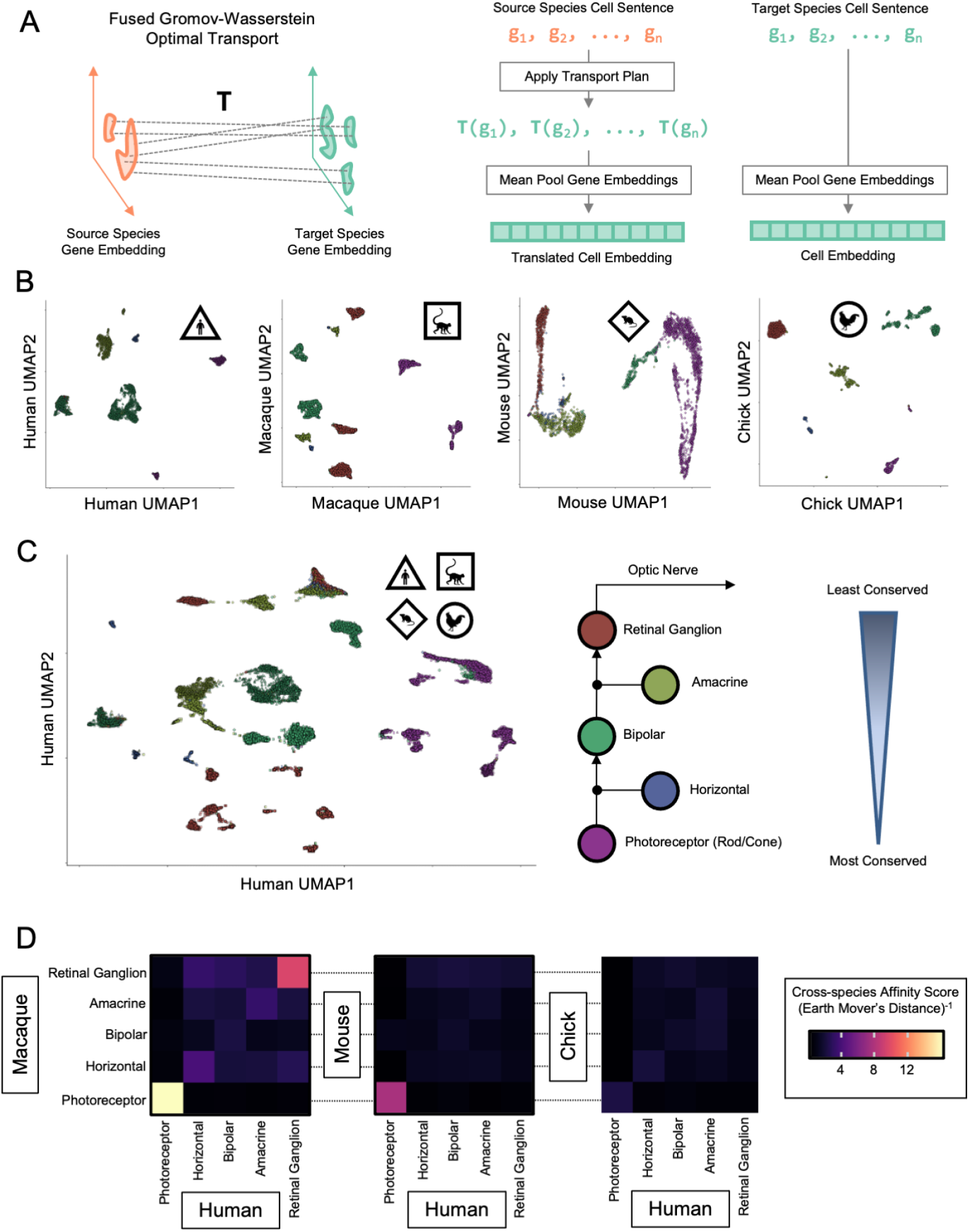
Cross-species translation and integration of retinal neurons reveals patterns of evolutionary conservation. (A) Fused Gromov-Wasserstein optimal transport can map gene embeddings across species, and gene embeddings can be pooled to generate cell sentence embeddings. (B) Cell sentence embeddings generated using word2vec are concordant with published expert-defined cell type labels. (C) Major retinal cell types cluster coherently across species after cell sentence translation and unified embedding. (D) Inverse earth mover’s distance shows decreasing cross-species affinity with increasing evolutionary distance.

Leveraging the cell sentence representation, we formulated the problem of mapping cell expression profiles across species as one of translation. Differences in genes across species can be thought of as differences in *vocabulary*, while differences in the interactions between genes may be though of as differences in *grammar*. While many traditional approaches to automated machine translation rely on the availability of matched bilingual corpora (pairs of sentences with equivalent meaning in both languages), several approaches have been recently developed allowing translation between languages using monolingual data alone.

One such approach, which we have adapted for cross-species translation, applies Gromov optimal transport to match word embeddings across languages [20]. Our adapted approach also incorporates prior knowledge of gene homologs by using fused Gromov-Wasserstein optimal transport, which smoothly interpolates between pure Wasserstein (minimizing feature distances) and pure Gromov (minimizing structural distances) optimal transport, with cost weighting subject to a hyperparameter [21](Fig 2A). Using this approach, we mapped cells from the five major retinal neuronal cell types from rhesus macaque, mouse, and chick onto the human gene embedding space, allowing for integrated analysis (Fig 2C). In the integrated representation, cells of the same type clustered together across different species (Fig 2C). The distances between cells increased with evolutionary distance, as well as with increasing laminar distance from the primary sensory neurons (photoreceptors), consistent with previous reports [16] (Fig 2D).

One potential limitation of our method is that the transformation of cell transcript count data to cell sentences results is a lossy conversion, as only per-cell relative ordering information is retained, and not absolute counts. This means that order-preserving changes to gene expression will not change in the cell sentence representation. Conversely, these properties may also make cell sentence representations more robust to batch-based variation in sequencing depth, or mRNA recovery during library preparation.

Our experiments with cell2sentence demonstrate that abundance-ordering is a simple and robust data transformation that permits the analysis of single-cell transcriptomics data sets using tools from natural language processing. Cross-species translation in particular represents a promising area that may further our understanding of human disease models in other organisms and promote the development of new therapeutics targeting conserved mechanisms.

## 2 Methods

### 2.1 Notation and Definitions

We consider a cell by gene count matrix ***C*** as a non-negative integer-valued matrix with *n* rows and *k*, and *C_i,j_* is the number of molecules detected for gene *j* in cell *i*. We define the cell sentence *s_i_* as the ordered list of non-zero gene in cell ***C***_*i*,_:, sorted in order of decreasing gene abundance, with ties broken at random. We define a function *S* mapping from a count matrix ***C*** to the corresponding set of cell sentences 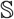, such that 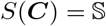.

In all experiments, only the top 2,000 highly variable genes were included in cell sentence generation, calculated as per the `seurat_v3` option for highly variable gene calculation in the `scanpy` library.

### 2.2 Edit distances defined between cell sentences

Levenshtein, Damerau–Levenshtein, Jaro, and Jaro-Winkler edit distances were defined in the usual way and calculated using the jellyfish python library. To increase distance matrix calculation speed, only the first 20 genes (prefix length 20) in each cell sentence were considered in the edit distance calculation. The zlib-normalized compression distance (zlib-NCD) was calculated as:

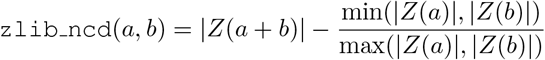

Where *Z*() is the zlib utility from python with default compression parameters, | · | denotes the byte length of the enclosed expression, and *a* and *b* are UTF-8 encoded byte string representations of the cell sentences.

### 2.3 Gene embeddings trained from cell sentences

By rendering the cell sentence representation as text, where gene names correspond to words, a gene embedding was generated using the standard published word2vec implementation for retinal scRNAseq data sets from human (*Homo sapiens*), rhesus macaque (*Macaca mulatta*), mouse (*Mus musculus*), and chicken (*Gallus gallus*) [15, 16, 17, 18]. Embeddings were trained on all retinal cells from the provided datasets without filtering. Word2vec was run using the continuous bag of words (CBOW) objective, with a window size of 8, 15 iterations, and an output embedding of 200 dimensions on all cell sentences concatenated into a single line.

The 50 ‘‘hallmark” gene sets from the molecular signatures database (MSigDB) were used to determine the average distance between genes within a gene set, as compared to between gene sets [19]. For each hallmark gene set with greater than 5 genes for which an embedding was available, 100 pairs of genes were selected at random where both members were in the set (Within Set), and another 100 pairs for which only a single member was in the set (Between Set).

### 2.4 Cell sentence embeddings from gene embeddings

Given a gene *g*, we define its *k* dimensional embedding as *ĝ* ∈ ℝ^*k*^. We may define the cell embedding for a cell sentence *s* = {*g*_1_, *g*_2_,…, *g_n_*} as:

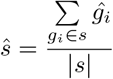

Noting that no positional information is incorporated into the cell sentence embedding, only gene presence or absence.

### 2.5 Fused gromov-wasserstein optimal transport

The implementation for fused Gromov-Wasserstein optimal transport from the python `pot` library [21]. Default parameters were used, with uniform density used for the source and target transport distributions. A default tradeoff parameter of *α* = 0.5 was used and the metric cost matrix ***M*** for features across domains was defined as:

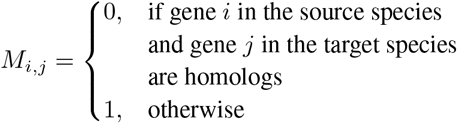

Fused Gromov-Wasserstein optimal transport yields a transport matrix ***T*** ∈ ℝ^*n*×*m*^ where *n* is the number of possible genes in the source language (i.e. the source language vocabulary size) and *m* is the number of possible genes in the target language.

A translated cell embedding for a cell sentence *s* in the source language is defined by mapping the genes from the source to the target language with the transport matrix ***T*** and then applying the weighted gene embeddings in the target language. Considering 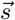 as a membership vector where

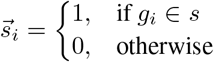

and ***E***_*target*_ is an embedding matrix in the target language where the *i*th row corresponds to the embedding vector for the *i*th gene, we may define the translated cell embedding for *s* in the target language as

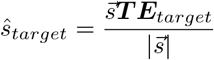

### 2.6 Statistics and visualizations

All distances between embedding vectors were calculated using the cosine distance implementation from the `scipy` library [22]. Significance for hallmark gene set coherence within the PCA, UMAP, and word2ec gene embedding spaces tested was assessed by the paired Wilcoxon signed-rank test.

For plots of cell embeddings, 10% of the cell sentences were randomly selected from each of the retinal data sets for evaluation. Plots were generated using the `plotnine` library for python as well as the `ggplot2` library for R [23]. In the heatmap visualization for cross-species translation, per-celltype affinities were defined as the multiplicative inverse of the earth mover’s distance between the two cell populations in the human gene embedding space. Figure graphics were created using resources from Flaticon.com.

## Data availability

All data used in the manuscript are publicly available. PBMC data are available from 10x Genomics at https://www.10xgenomics.com/resources/datasets. Human retinal data was obtained from GEo under accession GsE148077. Macaque single cell data was obtained from the Broad single Cell portal under study accession number sCP212. Mouse developmental retina data was obtained from GEo under accession GsE118614. Chick retina data was also obtained from GEo under accession GsE159107.

## Code availability

Code for cell2sentence is available in a public GitHub repository under the Creative Commons Attribution-Noncommercial-ShareAlike 4.0 License at: https://github.com/rahuldhodapkar/cell2sentence.

## Notes

https://github.com/rahuldhodapkar/cell2sentence

### Competing Interest Statement

The authors have declared no competing interest.

